# A robust SARS-CoV-2 replication model in primary human epithelial cells at the air liquid interface to assess antiviral agents

**DOI:** 10.1101/2021.03.25.436907

**Authors:** Thuc Nguyen Dan Do, Kim Donckers, Laura Vangeel, Arnab K. Chatterjee, Philippe A. Gallay, Michael D. Bobardt, John P. Bilello, Tomas Cihlar, Steven De Jonghe, Johan Neyts, Dirk Jochmans

**Affiliations:** KU Leuven - Department of Microbiology, Immunology and Transplantation, Rega Institute, Laboratory of Virology and Chemotherapy, Leuven, Belgium; CALIBR - Department of Medicinal Chemistry, the Scripps Research Institute, La Jolla, CA, USA; CALIBR - Department of Immunology and Microbial Science, the Scripps Research Institute, La Jolla, CA, USA; Gilead Sciences, Inc., Foster City, CA, USA

**Keywords:** SARS-CoV-2, antivirals, primary human airway epithelial cells, HAEC, remdesivir, GS-441524, EIDD-1931, AT-511, IFN

## Abstract

There are, besides remdesivir, no approved antivirals for the treatment of SARS-CoV-2 infections. To aid in the search for antivirals against this virus, we explored the use of human tracheal airway epithelial cells (HtAEC) and human small airway epithelial cells (HsAEC) grown at the air/liquid interface (ALI). These cultures were infected at the apical side with one of two different SARS-CoV-2 isolates. Each virus was shown to replicate to high titers for extended periods of time (at least 8 days) and, in particular an isolate with the D614G in the spike (S) protein did so more efficiently at 35°C than 37°C. The effect of a selected panel of reference drugs that were added to the culture medium at the basolateral side of the system was explored. Remdesivir, GS-441524 (the parent nucleoside of remdesivir), EIDD-1931 (the parent nucleoside of molnupiravir) and IFN (β1 and λ1) all resulted in dose-dependent inhibition of viral RNA and infectious virus titers collected at the apical side. However, AT-511 (the free base form of AT-527 currently in clinical testing) failed to inhibit viral replication in these *in vitro* primary cell models. Together, these results provide a reference for further studies aimed at selecting SARS-CoV-2 inhibitors for further preclinical and clinical development.

## INTRODUCTION

Remdesivir is currently the only approved antiviral for the treatment of COVID-19^1^. Major efforts are ongoing to develop novel antiviral drugs for effective COVID-19 treatment. To aid in their development, physiologically-relevant *in vitro* models are needed. Immortalized cell lines originating from non-respiratory (and often non-human) tissues are frequently used in early preclinical studies for antiviral assessment. For example, VeroE6, a widely used cell line in SARS-CoV-2 studies, is defective in the expression of main SARS-CoV-2 receptors (angiotensin-converting enzyme 2 (ACE2) and transmembrane protease serine 2 (TMPRSS2)). Hence, screening campaigns often result in the discovery of antiviral agents that regulate autophagy pathways and endosomal-lysosomal maturation, which may not be pertinent or translatable as SARS-CoV-2 therapies^2^. Meanwhile, air-liquid interface (ALI) of differentiated primary human airway epithelial cells (HAEC) possess the architecture and cellular complexity of human lung tissue and are permissive to variety of respiratory viral infections^3,4^. Containing all relevant cell types of the lower respiratory tract (ciliated, goblet and basal cells), which includes ACE2 and TMPRSS2 expressing cells, this system allows dissection of the host-pathogen interactions at molecular and cellular levels and provides a platform for profiling antiviral drugs.

In this study, we explored the effects of several reported SARS-CoV-2 inhibitors on the replication of different SARS-CoV-2 isolates in HAEC ALI cultures focusing the testing primarily on the nucleoside class of inhibitors with previously reported in vitro anti-SARS-CoV-2 activity. Our results provide a reference set of data for these relevant in vitro cell models to support the preclinical development of SARS-CoV-2 inhibitors.

## MATERIALS AND METHODS

### Cells and virus isolates

The African monkey kidney cell line VeroE6 tagged green fluorescent protein (VeroE6-GFP, kindly provided by M. van Loock, Janssen Pharmaceutica, Beerse, Belgium) and VeroE6 were maintained in Dulbecco’s modified Eagle’s medium (DMEM; Gibco, catalogue no. 41965-039) supplemented with 10% v/v heat-inactivated foetal bovine serum (HI-FBS; HyClone, catalogue no. SV03160.03), 1% v/v sodium bicarbonate 7.5% w/v (NaHCO_3_; Gibco, catalogue no. 25080-060), and 1% v/v Penicillin-Streptomycin 10000 U/mL (P/S; Gibco, catalogue no. 15140148) at 37°C and 5% CO_2_. The hepatocellular carcinoma cell line Huh7 (kindly provided by Ralf Bartenschlager, University of Heidelberg, Germany) was propagated in DMEM supplemented with 10% HI-FBS, 1% NaHCO_3_, 1% P/S, 1% non-essential amino acids (NEAA; Gibco, catalogue no. 11140050), and 2% HEPES 1M (Gibco, catalogue no. 15630106) at 37°C and 5% CO_2_. All assays involving virus growth were performed in the respective cell growth medium containing 2% (VeroE6-GFP) or 4% (Huh7) instead of 10% FBS.

SARS-CoV-2 isolate BetaCoV/Germany/BavPat1/2020 (EPI_ISL_406862|2020-01-28, kindly provided by C. Drosten, Charité, Berlin, Germany) and BetaCov/Belgium/GHB-03021/2020 (EPI_ISL_407976|2020-02-03) retrieved from RT-qPCR-confirmed COVID-19 positive patients in January and February 2020 were described previously^5,6^. The generation of virus stocks by serial passaging in Huh7 and VeroE6 cells were fully reported^7,8^. BavPat1 isolate (passage 2 (P2)) and GHB-03021 isolate (P6 and P7) were used for the air liquid-interface experiment while only the latter was used for standard *in vitro* assays in VeroE6-GFP cells (P6 and P7) and in Huh7 cells (P9). The genomic sequence of both isolates is highly similar. BavPat1 carries the D614G amino acid change in the spike-protein while the GHB-03021 has a ΔTQTNS deletion at 676-680 residues that is typical for SARS2 strains that have been passaged several times on VeroE6 cells. All infectious virus-containing works were conducted in biosafety level 3 (BSL-3) and 3+ (CAPs-IT) facilities at the Rega Institute for Medical Research, KU Leuven, according to institutional guidelines.

### Compounds

Remdesivir was synthesized at Gilead Sciences, Inc. (Foster City, CA). GS-441524 was either synthesized at Gilead (**Figure 2**) or it was purchased from Carbosynth (United Kingdom) (other figures). EIDD-1931 was purchased from R&D Systems (USA). Stock solutions (10 mM) were prepared using analytical grade dimethyl sulfoxide (DMSO). AT-511 was synthesized and chemically validated at the California Institute for Biochemical Research (Calibr) (La Jolla, CA) and used as a 10 mM DMSO solution. The biological activity of AT-511 was confirmed in an antiviral assay with hepatitis C virus (data not shown). IFN λ1 was purchased from R&D Systems and IFN β-1a was a kind gift from the laboratory of Immunobiology (Rega Institute, KU Leuven, Belgium). Both were reconstituted in sterile phosphate buffered saline (PBS, Life Technologies) containing at least 0.1% FBS.

### *In vitro* standard antiviral and toxicity assays

VeroE6-GFP cells were seeded at a density of 25000 cells/well in 96-well plates (Greiner Bio One, catalogue no. 655090) and pre-treated with three-fold serial dilutions of the compounds overnight. On the next day (day 0), cells were infected with the SARS-CoV-2 inoculum at a multiplicity of infection (MOI) of 0.001 median tissue infectious dose (TCID50) per cell. The number of fluorescent pixels of GFP signal determined by High-Content Imaging (HCI) on day 4 post-infection (p.i.) was used as a read-out. Percentage of inhibition was calculated by subtracting background (number of fluorescent pixels in the untreated-infected control wells) and normalizing to the untreated-uninfected control wells (also background subtracted). The 50% effective concentration (EC_50_, the concentration of compound required for fifty percent recovery of cell-induced fluorescence) was determined using logarithmic interpolation.

Potential toxicity of compounds was assessed in a similar set-up in treated-uninfected cultures where metabolic activity was quantified at day 5 using the MTS assay as described earlier^9^. The 50% cytotoxic concentration (CC_50_, the concentration at which cell viability reduced to 50%) was calculated by logarithmic interpolation.

Huh7 cells were pre-seeded at 6000 cells/well in 96 well-plates (Corning, catalogue no.3300) and incubated overnight at 37°C and 5% CO_2_. On day 0, cells were firstly treated with the three-fold serial dilution of a potential antiviral, followed by either the inoculation of SARS-CoV-2 at MOI of 0.0037 TCID50/cell or addition of fresh medium. After 4 days, differences in cell viability caused by virus-induced cytopathic effect (CPE) or by compound-specific toxicity were evaluated using MTS assays. The EC_50_ and CC_50_ were calculated as above-mentioned.

### Viral infection of reconstituted human airway epithelium cells

Human tracheal airway epithelium cells (HtAEC; catalogue no. EP01MD) and human small airway epithelium cells (HsAEC; catalogue no. EP21SA) from healthy donors were obtained from Epithelix (Geneva, Switzerland) in an air-liquid interphase set-up. After arrival, the insert was washed with pre-warmed 1x PBS (Gibco, catalogue no. 14190-094) and maintained in corresponding MucilAir medium (Epithelix, catalogue no. EP04MM) or SmallAir medium (Epithelix, catalogue no. EP64SA) at 37°C and 5% CO_2_ for at least 4 days before use. On the day of the experiment, the HAEC were first pre-treated with basal medium containing compounds at different concentrations for indicated hours, followed by exposing to 100 µL of SARS-CoV-2 inoculum from apical side for 1.5 hours. Then the cultures were incubated at the indicated temperatures. The first apical wash with PBS was collected either right after the removal of viral inoculum (day 0) or 24 hours later (day 1 post-infection (p.i.)). Every other day from day 0, subsequent apical washes were collected whereas compound-containing medium in the basolateral side of the HAEC culture was refreshed. Wash fluid was stored at -80°C for following experiments.

### RNA extraction and quantitative reverse transcription-PCR (RT-qPCR)

Viral RNA in the apical wash was isolated using the Cells-to-cDNA™ II cell lysis buffer kit (Thermo Fisher Scientific, catalogue no. AM8723). Briefly, 5 µL wash fluid was added to 50 µL lysis buffer, incubated at room temperature (RT) for 10 min and then at 75°C for 15 min. 150 µL nuclease-free water was additionally added to the mixture prior to RT-qPCR. In parallel, a ten-fold serial dilution of corresponding virus stock was extracted. The amount of viral RNA expressed as TCID50 equivalent per insert (TCID50e/insert) was quantified by RT-qPCR using iTaq universal probes one-step kit (Bio-Rad, catalogue no. 1725141), and a commercial mix of primers for N gene (forward primer 5’-GACCCCAAAATCAGCGAAAT-3’, reverse primer 5’-TCTGGTTACTGCCAGTTGAATCTG-3’) and probes (5’-FAM-ACCCCGCATTACGTTTGGTGGACC-BHQ1-3’) manufactured at IDT Technologies (catalogue no. 10006606). The reaction (final volume: 20 µL) consisted of 10 µL one-step reaction mix 2X, 0.5 µL reverse transcriptase, 1.5 µL of primers and probes mix, 4 µL nuclease-free water, and 4 µL viral RNA. The RT-qPCR was executed on a Lightcycler 96 thermocycler (Roche), starting at 50°C for 15 min and 95°C for 2 min, followed by 45 cycles of 3 sec at 95°C and 30 sec at 55°C.

### Titration using a 50% tissue culture infectious dose (TCID50) assay

VeroE6 cells were seeded in 96-well tissue culture plates (Corning, catalogue no. 353072) at a density of 1×10^4^ cells/180 µL/well. After 24 hours, serial 10-fold dilutions of ALI wash fluid were prepared in the plates. Cells were incubated for 3 days at 37°C and evaluated microscopically for the absence or presence of virus induced cytopathic effect (CPE). The infectious viral titer was determined by end-point titration, expressed as TCID50/ml. Virus titers were calculated by using the Spearman and Karber method as previously reported^10,11^.

### Statistical analysis

All statistical comparisons in the study were performed in GraphPad Prism (GraphPad Software, Inc.). Statistical significance was determined using the ordinary one-way ANOVA with Dunnett’s multiple comparison test. P-values of ≤0.05 were considered statistically significant. In the figures “ns” indicates a p > 0.05. Asterisks indicate a statistical significance level of *p < 0.05, **p < 0.01, ***p < 0.001, ****p < 0.0001.

## RESULTS

### Replication kinetics of SARS-CoV-2 in reconstituted human airway epithelia

We first compared the replication kinetics of the Belgian isolate GHB-03021 and the German isolate BavPat1 of SARS-CoV-2. The main differences in the genomes of these viruses is the D614G amino acid change in the spike-protein of BavPat1 and the deletion of several amino acids near the furin-cleavage site in the GHB-03021 isolate (because of extensive passaging in VeroE6 cells). The replication kinetics was investigated at respectively 35 and 37°C in both cultures from tracheal cells (HtAEC) or from small airway cells (HsAEC). In preliminary experiments, it was observed that an input of 10^2^, 10^3^ or 10^4^ TCID50/insert resulted in comparable levels of virus production (data not shown). We therefore selected 2×10^3^ TCID50/insert as the viral input for this experiment. Overall, BavPat1 infected the cultures more efficiently than the GHB-03021 isolate did (**Figure 1**). For example, at 37°C not all bronchiole-airway derived inserts infected with GHB-03021 resulted in a productive infection whereas all cultures infected with BavPat1 showed productive infection under all conditions. Virus replication was for both isolates higher at 35°C and more reproducible as compared with a temperature of 37°C. This difference was more pronounced for the HsAEC than for the HtAEC cultures.

**Figure 1.**
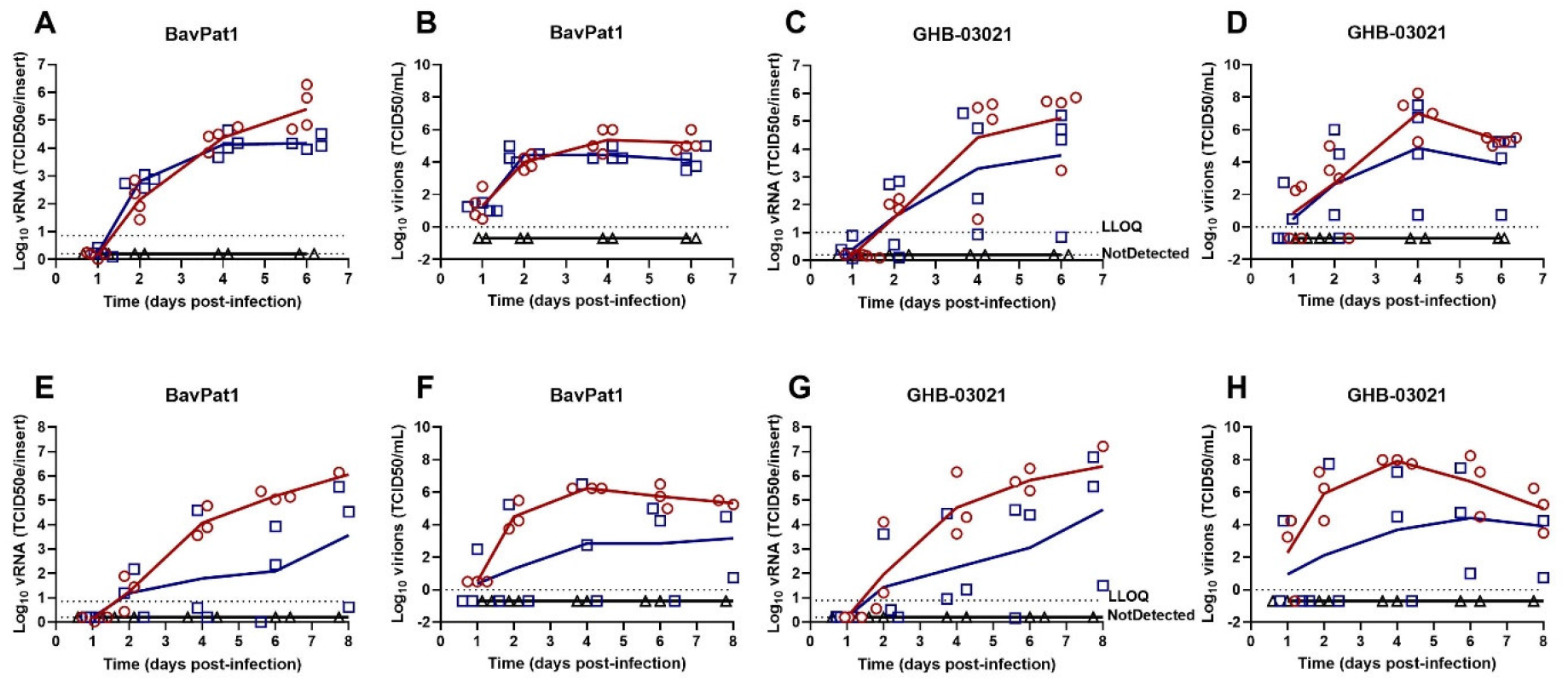
SARS-CoV-2 replication kinetics in air-liquid interface cultures. Viral replication of BavPat1 and GHB-03021 isolates at 2×10^3^ TCID50/insert in human tracheal airway epithelia cells (A-D) or in human small airway epithelia cells (E-H) at either 35°C (red circle) or 37°C (blue square) in comparison with uninfected control (black triangle). Viral RNA or infectious particles in apical washes were quantified by RT-qPCR (A, C, E, G) or by end-point titrations (B, D, F, H), respectively. The results of individual inserts are depicted as dots. The line follows the average of each conditions.

**Figure 2.**
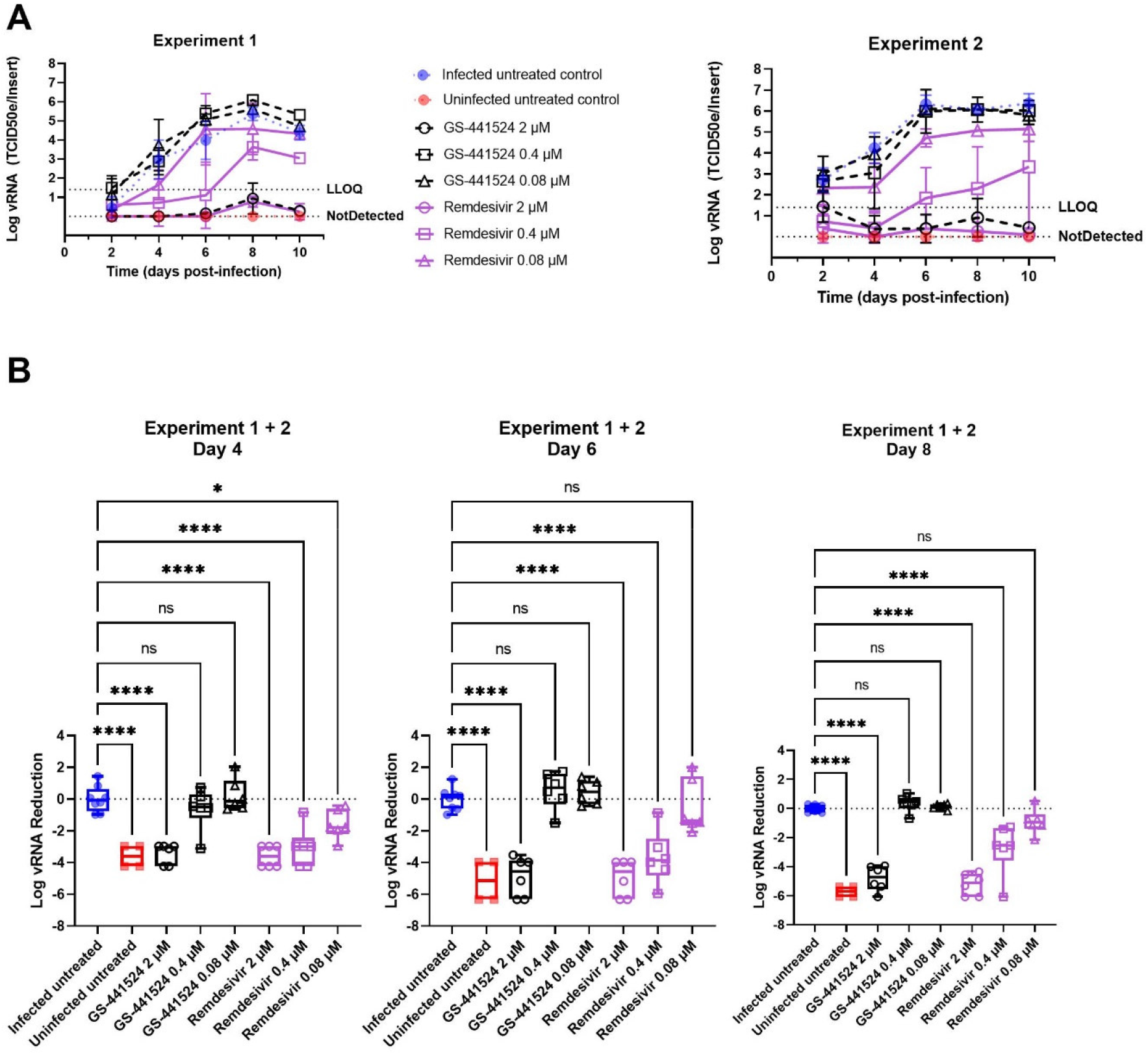
Remdesivir and GS-441524 inhibit SARS-CoV-2 replication in human tracheal airway cultures. Compounds were added to the basal medium starting 2 h before infection and treatment continued for 10 days. HtAEC cultures were infected with SARS-CoV-2 GHB-03021 at 2×10^4^ TCID50/insert and incubated at 37°C. Viral RNA in apical washes was quantified by RT-qPCR. (A) Kinetics of viral replication. Two independent experiments with HtAEC from different donors were performed. In each experiment 3 independent inserts were used for each compound concentration. LLOQ represent the lower limit of quantification. (B) The Log vRNA Reduction on day 4, 6 and 8 of the experiment. For this analysis the samples of both experiments were pooled. The Log vRNA Reduction was calculated by subtracting the average Log vRNA of the infected untreated samples for that particular day from the Log vRNA of each sample. ns p > 0.05, *p < 0.05, **p < 0.01, ***p < 0.001, ****p < 0.0001.

### Effect of selected antivirals on SARS-CoV-2 replication in HAEC cultures

Four nucleoside analogues that are known inhibitors of SARS-CoV-2 replication were selected as reference for studies in HAEC cultures. These included remdesivir, GS-441524^12-16^ (the parent nucleoside of remdesivir), EIDD-1931^16-18^ (the parent nucleoside of molnupiravir) and AT-511^17^ [a guanosine nucleotide analogue with activity against hepatitis C virus (HCV)]. To select a suitable concentration range of these molecules, we first explored their effect in VeroE6 and Huh7 cell lines. Treatment with remdesivir, GS-441524 and EIDD-1931 selectively inhibited SARS-CoV-2 GHB-03021 replication. On the other hand, AT-511 was entirely devoid of antiviral activity (**Table 1**).

**Table 1.**
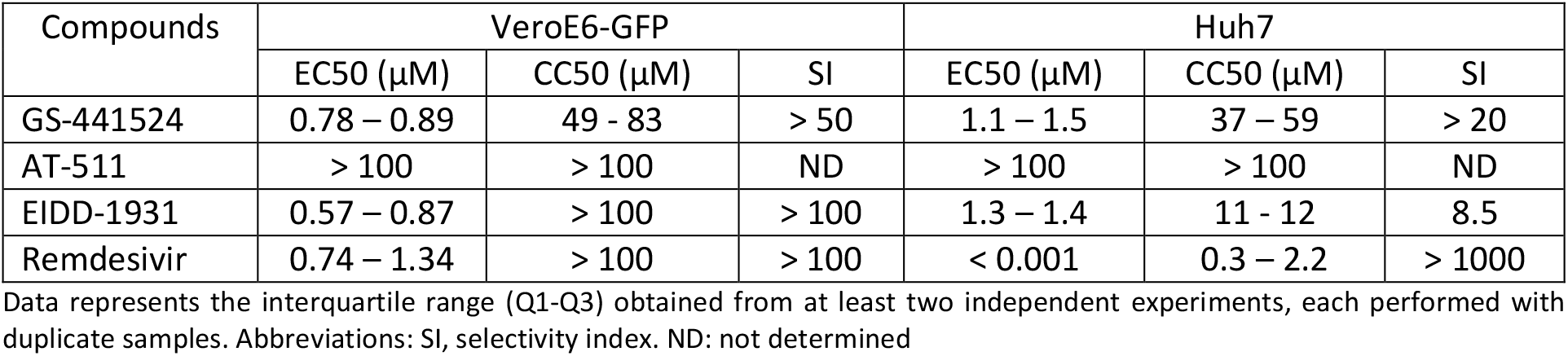
Antiviral activity against SARS-CoV-2 in cell culture.

| Compounds | VeroE6-GFP |  |  | Huh7 |  |  |
| --- | --- | --- | --- | --- | --- | --- |
|  | EC50 (μM) | CC50 (μM) | SI | EC50 (μM) | CC50 (μM) | SI |
| GS-441524 | 0.78 – 0.89 | 49 - 83 | > 50 | 1.1 – 1.5 | 37 – 59 | > 20 |
| AT-511 | > 100 | > 100 | ND | > 100 | > 100 | ND |
| EIDD-1931 | 0.57 – 0.87 | > 100 | > 100 | 1.3 – 1.4 | 11 - 12 | 8.5 |
| Remdesivir | 0.74 – 1.34 | > 100 | > 100 | < 0.001 | 0.3 – 2.2 | > 1000 |

Data represents the interquartile range (Q1-Q3) obtained from at least two independent experiments, each performed with duplicate samples. Abbreviations: SI, selectivity index. ND: not determined

In a subsequent experiment, the activities of a concentration range of remdesivir and GS-441524 were tested on SARS-CoV-2 GHB-03021 in HtAEC cultures during 10 days of treatment (**Figure 2**). Both compounds showed strong inhibition at 2 µM throughout the experiment, suppressing virus replication below the limit of quantification. Remdesivir also showed inhibition at 0.4 µM and at 0.08 µM until day 4. However, while remdesivir was still highly potent at 0.4 µM (2 log_10_ reduction in vRNA), a steady increase in virus replication over time was observed at this concentration.

Next we explored the activity of GS-441524 in HtAEC cultures infected with SARS-CoV-2 GHB-03021 and in HsAEC cultures infected with BavPat1 (**Figure 3**). At 10 µM, GS-441524 sterilized the HtAEC cultures from the GHB-03021 virus. Indeed, no virus production was detected during the first 9 days of treatment and, when treatment was stopped, no rebound was observed over the next 5 days of culturing. When evaluated at a concentration of 1 µM, GS-441524 reduced virus yield by ∼1 log_10_ during the time of treatment, but lost activity once the compound was removed from the culture. In a separate experiment, GS-441524 at 3 µM resulted in complete inhibition of virus production upon infection with BavPat1 (**Figure 3F-H**). Also, 10 µM of EIDD-1931 resulted in a pronounced antiviral effect (**Figure 4F-J**), whereas AT-511 was surprisingly devoid of any antiviral activity (**Figure 4A-E**) at the different concentrations tested (1 and 10 µM).

**Figure 3.**
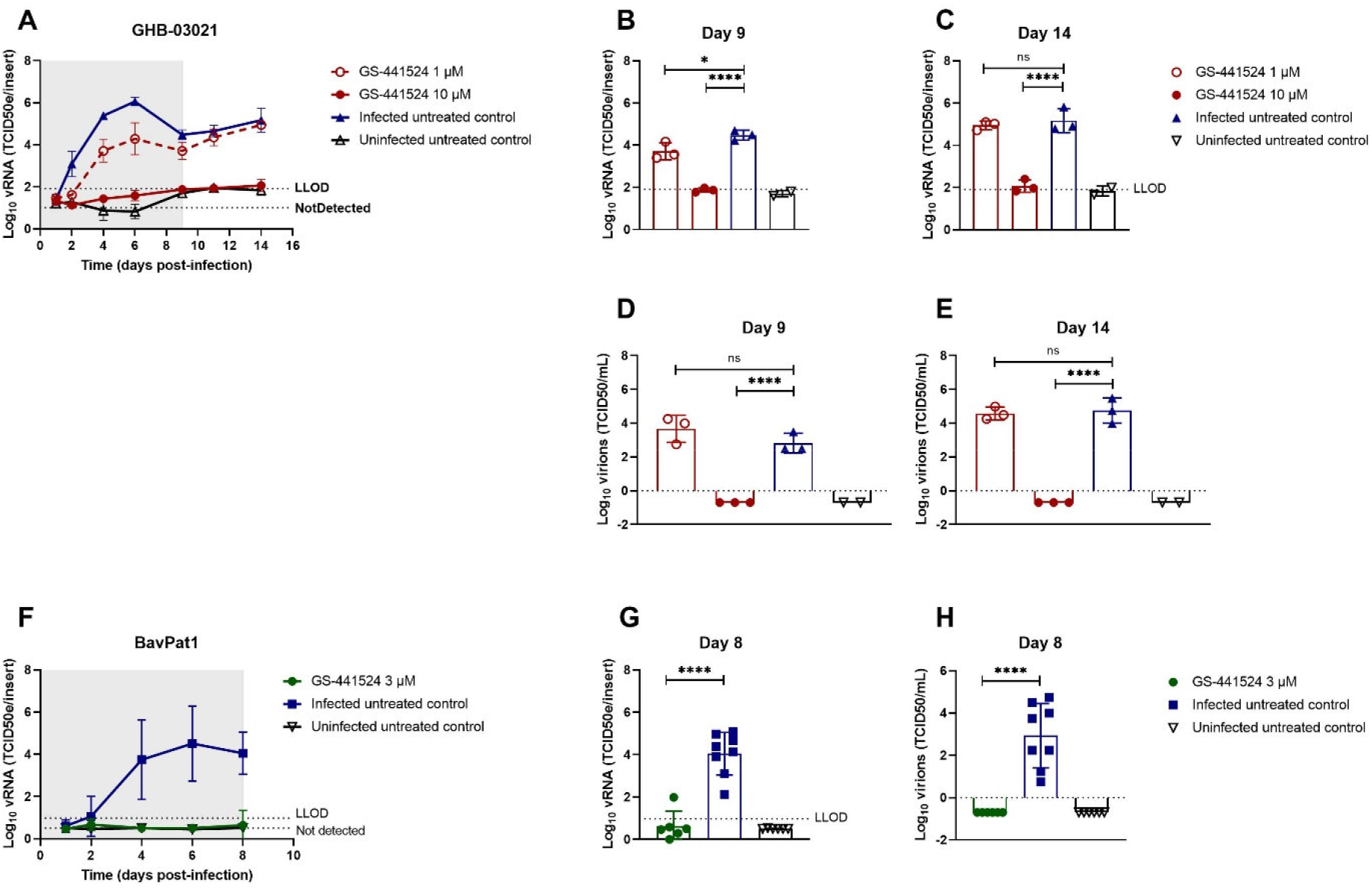
GS-441524 blocks SARS-CoV-2 replication in human tracheal airway cultures and human small airway cultures. Compounds were added to the basal medium at different concentrations, starting from the in vitro EC50. The treatment period is illustrated in grey, initiating at 2 hours before infection. Viral RNA or infectious particles in apical washes were quantified by RT-qPCR (A-C, F-G) or by end-point titrations (D-E, H), respectively. (A) Effect of GS-441524 at 10 and 1 µM on the replication of the GHB-03021 isolate. Tracheal issues were infected with 2×10^4^ TCID50/insert at 37°C. The viral production was compared between groups after 9 (B, D) or 14 days post-infection (C, E). (F) Activity of GS-441524 at 3 µM on the replication of the BavPat1 isolate. Small airway tissues were infected with BavPat1 isolate at 2×10^3^ TCID50/insert at 37°C. Viral production was compared between groups at the end of the treatment (G-H). All data are mean ± SD of at least three independent ALI inserts. The dotted lines represent the lower limit of detection (LLOD). Asterisks indicate a significant difference between treated samples and infected untreated control. *p < 0.05, **p < 0.01, ***p < 0.001, ****p < 0.0001

**Figure 4.**
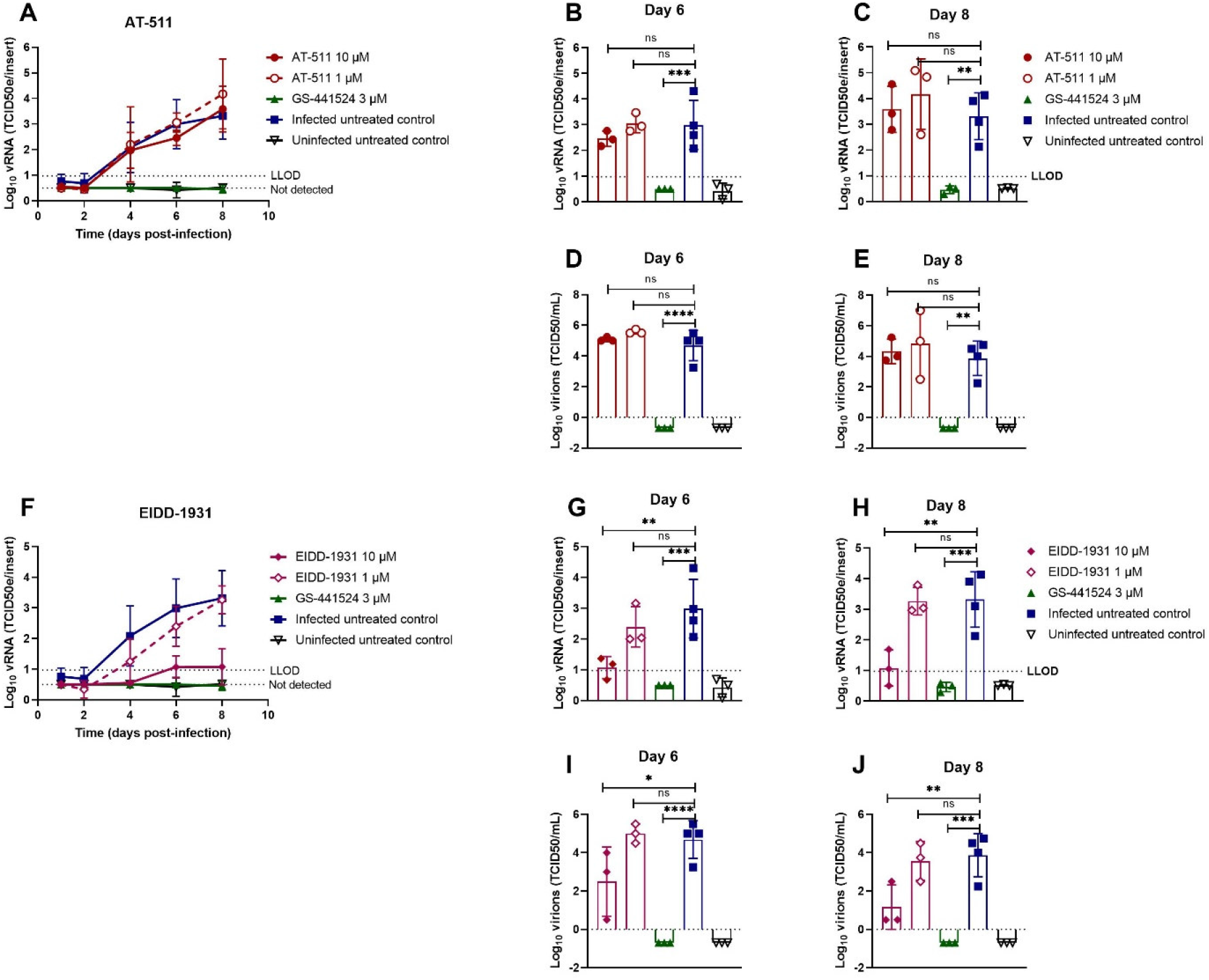
Assessment of antiviral activity of AT-511 and EIDD-1931 against SARS-CoV-2 in primary human tracheal airway cultures. Compounds were added to the basal medium at different concentrations, starting 1 hour before infection with BavPat1 isolate at 2×10^3^ TCID50/insert at 37°C. Basal medium, with or without compounds, was refreshed every other day from day 0 to day 8. Viral RNA or infectious particles in apical washes were quantified by RT-qPCR (A-C, F-H) or by end-point titrations (D-E, I-J), respectively. Dose-response and time-dependent activity of AT-511 (A) and EIDD-1931 (F). Comparison of the viral production between groups after 6 (B, D, G, I) or 8 days post-infection (C, E, H, J). All data are mean ± SD of at least three replicates. The dotted lines represent the lower limit of detection (LLOD). Asterisks indicate a significant difference between treated samples and infected untreated control. *p < 0.05, **p < 0.01, ***p < 0.001, ****p < 0.0001

### Prophylactic interferon type I and type III reduce SARS-CoV-2 production

Human IFN has previously been used to treat several viral infections^19,20^ and clinical trials against SARS-CoV-2 are ongoing (ClinicalTrials.gov number: NCT04315948, NCT04385095, and NCT04492475). We investigated whether IFN β-1a and IFN λ1 exert antiviral activity when used as a prophylactic monotherapy. Tracheal cultures were pre-treated with either 5 and 50 ng/mL IFN λ1 (5 ng/mL is the average concentration secreted in the basal medium of infected HAEC cultures^21^) or 1 and 100 IU/mL IFN β-1a for 24 hours, and subsequently infected with BavPat1. Both drugs were able to reduce viral titers in a dose-dependent manner (**Figure 5A, 5F**). Viral loads were reduced by 100 IU/mL IFN β-1a (3.3 log_10_ vRNA reduction, 3.6 log_10_ titer reduction) and 50 ng/mL IFN λ1 (4.2 log_10_ vRNA reduction, 5.0 log_10_ titer reduction) on day 4 p.i. (**Figure 5B, 5D, 5G, 5I** respectively). At later time points viral load in the treated samples increased again.

**Figure 5.**
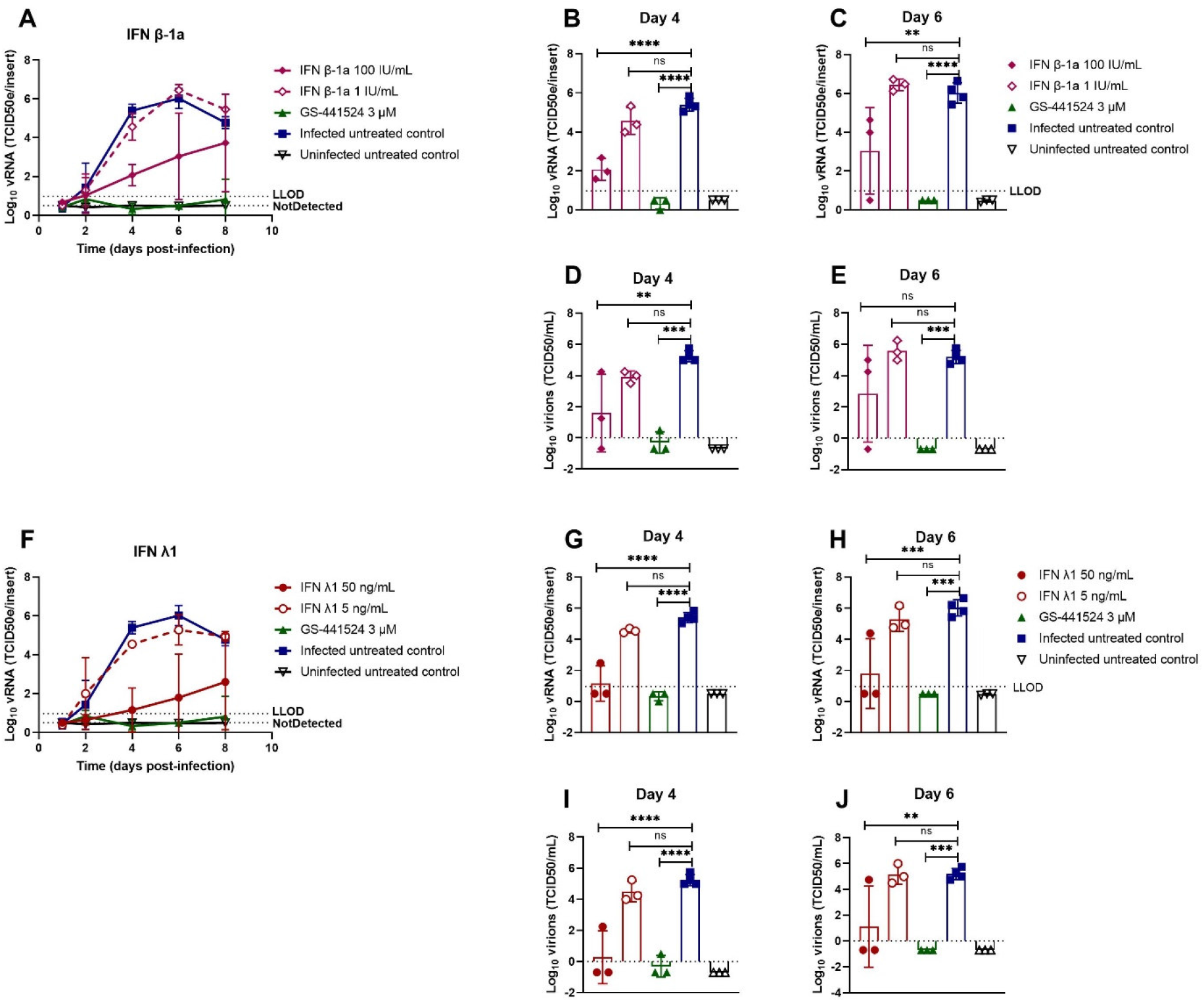
Prophylactic interferon type I and type III reduce SARS-CoV-2 production. IFNs were added to the basal medium at different concentrations 24 hours before infection with BavPat1 isolate (2×10^3^ TCID50/insert at 37°C) and medium with or without IFNs was refreshed every other day from day 0 to day 8. Viral RNA or infectious virions in apical washes were quantified by RT-qPCR (A-C, F-H) or by end-point titrations (D-E, I-J), respectively. Dose-response and time-dependent activity of AT-511 (A) and EIDD-1931 (F). The viral production was compared between groups after 4 (B, D, G, I) or 6 days post-infection (C, E, H, J). All data are mean ± SD of at least three replicates. The dotted lines represent the lower limit of detection (LLOD). Asterisks indicate a significant difference between treated samples and infected untreated control. *p < 0.05, **p < 0.01, ***p < 0.001, ****p < 0.0001

## DISCUSSION

We demonstrate that *ex vivo* models reconstituted from human tracheal or small airway epithelium are permissive for SARS-CoV-2 infection and robustly produces viral progenies from the apical side in long-term experiments (up to 14 days p.i.). Recent studies have also reported on the effect of different SARS-CoV-2 isolates and incubation temperatures on virus replication kinetics^22-25^. We tested two isolates, BavPat1 and GHB-03021, and found that BavPat1 proved to be more infectious. The BavPat1 isolate carries the p.D614G substitution in the spike (S) protein, which has been reported to increase the stability and infectivity of virions in HAEC culture by enhancing the ACE2-receptor-binding^26,27^. Isolates with this p.D614G substitution have become globally dominant^26-29^. Whereas the GHB-03021 has a deletion of several amino-acids in the S1/S2 boundary that is typically found in VeroE6-adapted isolates^7^. Continued propagation of SARS-CoV-2 in Vero cells are reported to cause several substitutions and/or deletions in the S1/S2 boundary^25,30-33^, which are only rarely observed in clinical samples^31,32^. We speculate that the adaptation to Vero cells results in a phenotype that allows more efficient entry through an ACE2-independent pathway. This entry mechanism would enhance entry in VeroE6 cells but would limit entry in primary lung epithelial cells. Further mechanistic studies are required to elucidate this hypothesis.

The anatomical distance and ambient temperature between upper and lower human respiratory tracts have a profound influence on the replication kinetics of respiratory viruses^22,34-36^. In agreement with other studies, we observed SARS-CoV-2 growth in favour of lower temperature (35°C) which can be attributed to the temperature preference of SARS-CoV-2 S protein for its folding and transport^24,37^. Both primary HtAEC and HsAEC cultures are shown to be a robust model for SARS-CoV-2 replication that can be used for antiviral drug profiling.

A promising target for the development of novel antiviral agents active against coronaviruses is the viral RNA-dependent RNA polymerase (RdRp)^38^. Remdesivir, a phosphoramidate prodrug of an adenosine C-nucleoside, was approved as the first COVID-19 therapy for the treatment of hospitalized patients based on several randomized clinical studies. Our experiments showed that remdesivir had strong SARS-CoV-2 antiviral potency at 2 µM and intermediate potency at 0.4 µM for up to 10 days of treatment. Partial viral suppression persisted at 0.08 µM until 4 days p.i.. These results are in agreement with an earlier study^13^, where the virus replication was monitored for 3 days. We demonstrated that the parent nucleoside GS-441524^12-16^ at 2 µM can also “sterilize” HAEC cultures from SARS-CoV2 as no rebound of the virus was noted several days after removal of the molecule. At 1 µM GS-441524, an intermediate antiviral potency was observed, but significant antiviral activity was not observed at 0.4 µM in the HAEC cultures. Differences in antiviral potencies of remdesivir and GS-441524 have been reported depending on the cell lines used, which correlates with the formation of the biologically active (5’-triphosphate) metabolite^12,13^. The oral bioavailability of GS-441524 has been reported to be low in rats and monkey, but high in dog^39^, suggesting that oral delivery of GS-441524 in human is not easy to predict.

AT-527 is currently being evaluated in phase II clinical trials for COVID-19 (ClinicalTrials.gov). Anti-SARS-CoV-2 activity of AT-511, the free base form of AT-527, was not observed in VeroE6, Huh7 cells nor in the HAEC cultures. This is in contrast with a recent publication where sub-micromolar activities of AT-511 were observed in very similar assay systems^17^. Currently, there is no data to reconcile this discrepancy. One possibility is that small differences in the assay conditions may substantially influence the metabolism of AT-511 to its active form and consequently impact its antiviral activity. As AT-511 is a double prodrug requiring activation by multiple enzymatic processes, it may be more susceptible to differential experimental conditions. The implications of these observations for *in vivo* efficacy are not clear, but should be further investigated.

The effect of EIDD-1931, which is the parent nucleoside of the ester prodrug molnupiravir (EIDD-2801), was also investigated. EIDD-1931 has been reported to exert antiviral activity against various human coronaviruses and molnupiravir is currently in clinical trials for SARS-CoV-2^17,18^. Initial interim data from a phase II study provides first evidence for antiviral activity in COVID patients (https://www.croiconference.org/abstract/reduction-in-infectious-sars-cov-2-in-treatment-study-of-covid-19-with-molnupiravir/). Treatment with EIDD-1931 resulted in a pronounced antiviral effect in HtAEC cultures at 10 µM, which was then lost at 1 µM. A previous study reported potent antiviral activity of EIDD-1931 against SARS-CoV-2 in HAEC at 0.1 µM and at 2 days p.i.^18^. It remains to be investigated whether the longer incubation time (6 and 8 days p.i.) reduces the activity of EIDD-1931 similarly to other nucleoside analogues tested.

A significant inhibitory effect of IFN β-1a and IFN λ1 was also noted, at high concentrations and in particular during the first few days of treatment. At later time-points, viral replication increased in the treated cultures, suggesting that the virus can escape the initial effect of IFN. The effective concentration of IFN β-1a used in this study is comparable with a clinically achievable concentration and is in line with other reports^19,20,40^.

In conclusion, both the replication of two SARS-CoV-2 isolates in HAEC cultures and the antiviral effect of representative and clinically relevant nucleoside inhibitors were assessed, providing a reference for this physiologically relevant model and its effective use in testing additional inhibitors of SARS-CoV2 replication.

## Acknowledgements

We thank Birgit Voeten, Niels Cremers, Tina van Buyten and Thibault Francken for their excellent technical assistance. We also thank Piet Maes for kindly providing the SARS-CoV-2 GHB-03021/2020 isolate used in this study. This research was partially supported by Bill & Melinda Gates Foundation (BMGF) under grant number INV-006366. T.N.D.D received the fellowship from European Union’s Horizon 2020 research and innovation programme under Marie Sklodowska-Curie grant agreement No. 812673 (OrganoVIR project). Part of this research work was performed using the ‘Caps-It’ research infrastructure (project ZW13-02) that was financially supported by the Hercules Foundation and Rega Foundation, KU Leuven.

## Author contributions

J.N. and D.J. conceptualized and supervised the project. T.N.D.D, D.J., J.P.B., and T.C. designed the research. T.N.D.D. and K.D. performed the ALI-related experiments. T.N.D.D., L.V., and D.J. analysed data. A.J.C, P.A.G, and M.D.B. characterized AT-511 structure and tested its activity against HCV. T.N.D.D. wrote the first draft of the manuscript. D.J., S.D.J., L.V., and J.N. edited the manuscript. L.V., D.J. and J.N. acquired the funding.

## Reference

1. Beigel JH, Tomashek KM, Dodd LE, et al. Remdesivir for the Treatment of Covid-19 — Final Report. New England Journal of Medicine. 2020;383(19):1813–1826.

2. Murgolo N, Therien AG, Howell B, et al. SARS-CoV-2 tropism, entry, replication, and propagation: Considerations for drug discovery and development. PLoS Pathog. 2021;17(2):e1009225.

3. Boda B, Benaoudia S, Huang S, et al. Antiviral drug screening by assessing epithelial functions and innate immune responses in human 3D airway epithelium model. Antiviral Res. 2018;156:72–79.

4. Loo SL, Wark PAB, Esneau C, Nichol KS, Hsu AC, Bartlett NW. Human coronaviruses 229E and OC43 replicate and induce distinct antiviral responses in differentiated primary human bronchial epithelial cells. Am J Physiol Lung Cell Mol Physiol. 2020;319(6):L926–L931.

5. Bohmer MM, Buchholz U, Corman VM, et al. Investigation of a COVID-19 outbreak in Germany resulting from a single travel-associated primary case: a case series. Lancet Infect Dis. 2020;20(8):920–928.

6. Dellicour S, Durkin K, Hong SL, et al. A Phylodynamic Workflow to Rapidly Gain Insights into the Dispersal History and Dynamics of SARS-CoV-2 Lineages. Mol Biol Evol. 2020.

7. Boudewijns R, Thibaut HJ, Kaptein SJF, et al. STAT2 signaling restricts viral dissemination but drives severe pneumonia in SARS-CoV-2 infected hamsters. Nat Commun. 2020;11(1):5838.

8. Rut W, Groborz K, Zhang L, et al. SARS-CoV-2 M(pro) inhibitors and activity-based probes for patient-sample imaging. Nat Chem Biol. 2021;17(2):222–228.

9. Jochmans D, Leyssen P, Neyts J. A novel method for high-throughput screening to quantify antiviral activity against viruses that induce limited CPE. J Virol Methods. 2012;183(2):176–179.

10. Kärber G. Beitrag zur kollektiven Behandlung pharmakologischer Reihenversuche. Naunyn-Schmiedebergs Archiv für experimentelle Pathologie und Pharmakologie. 1931;162(4):480–483.

11. Spearman C. The method of ‘right and wrong cases’ (‘constant stimuli’) without Gauss’s formulae. British Journal of Psychology.2:227–242.

12. Li Y, Cao L, Li G, et al. Remdesivir Metabolite GS-441524 Effectively Inhibits SARS-CoV-2 Infection in Mouse Models. Journal of Medicinal Chemistry. 2021.

13. Pruijssers AJ, George AS, Schafer A, et al. Remdesivir Inhibits SARS-CoV-2 in Human Lung Cells and Chimeric SARS-CoV Expressing the SARS-CoV-2 RNA Polymerase in Mice. Cell Rep. 2020;32(3):107940.

14. Shi Y, Shuai L, Wen Z, et al. The Preclinical Inhibitor GS441524 in Combination with GC376 Efficaciously Inhibited the Proliferation of SARS-CoV-2 in the Mouse Respiratory Tract. bioRxiv. 2020:2020.2011.2012.380931.

15. Xie X, Muruato AE, Zhang X, et al. A nanoluciferase SARS-CoV-2 for rapid neutralization testing and screening of anti-infective drugs for COVID-19. Nat Commun. 2020;11(1):5214.

16. Zandi K, Amblard F, Musall K, et al. Repurposing Nucleoside Analogs for Human Coronaviruses. Antimicrob Agents Chemother. 2020;65(1).

17. Good SS, Westover J, Jung KH, et al. AT-527, a double prodrug of a guanosine nucleotide analog, is a potent inhibitor of SARS-CoV-2 in vitro and a promising oral antiviral for treatment of COVID-19. Antimicrob Agents Chemother. 2021.

18. Sheahan TP, Sims AC, Zhou S, et al. An orally bioavailable broad-spectrum antiviral inhibits SARS-CoV-2 in human airway epithelial cell cultures and multiple coronaviruses in mice. Sci Transl Med. 2020;12(541).

19. Lokugamage KG, Hage A, de Vries M, et al. Type I Interferon Susceptibility Distinguishes SARS-CoV-2 from SARS-CoV. J Virol. 2020;94(23).

20. Strayer DR, Dickey R, Carter WA. Sensitivity of SARS/MERS CoV to interferons and other drugs based on achievable serum concentrations in humans. Infect Disord Drug Targets. 2014;14(1):37–43.

21. Essaidi-Laziosi M, Brito F, Benaoudia S, et al. Propagation of respiratory viruses in human airway epithelia reveals persistent virus-specific signatures. J Allergy Clin Immunol. 2018;141(6):2074–2084.

22. Corman VM, Eckerle I, Memish ZA, et al. Link of a ubiquitous human coronavirus to dromedary camels. Proc Natl Acad Sci U S A. 2016;113(35):9864–9869.

23. Pohl MO, Busnadiego I, Kufner V, et al. SARS-CoV-2 Variants Reveal Features Critical for Replication in Primary Human Cells. bioRxiv. 2021:2020.2010.2022.350207.

24. V’kovski P, Gultom M, Kelly J, et al. Disparate temperature-dependent virus – host dynamics for SARS-CoV-2 and SARS-CoV in the human respiratory epithelium. bioRxiv. 2020:2020.2004.2027.062315.

25. Zhu Y, Feng F, Hu G, et al. The S1/S2 boundary of SARS-CoV-2 spike protein modulates cell entry pathways and transmission. bioRxiv. 2020:2020.2008.2025.266775.

26. Korber B, Fischer WM, Gnanakaran S, et al. Tracking Changes in SARS-CoV-2 Spike: Evidence that D614G Increases Infectivity of the COVID-19 Virus. Cell. 2020;182(4):812-827.e819.

27. Yurkovetskiy L, Wang X, Pascal KE, et al. Structural and Functional Analysis of the D614G SARS-CoV-2 Spike Protein Variant. Cell. 2020;183(3):739–751 e738.

28. Plante JA, Liu Y, Liu J, et al. Spike mutation D614G alters SARS-CoV-2 fitness. Nature. 2020.

29. Zhang L, Jackson CB, Mou H, et al. The D614G mutation in the SARS-CoV-2 spike protein reduces S1 shedding and increases infectivity. bioRxiv. 2020.

30. Davidson AD, Williamson MK, Lewis S, et al. Characterisation of the transcriptome and proteome of SARS-CoV-2 reveals a cell passage induced in-frame deletion of the furin-like cleavage site from the spike glycoprotein. Genome Medicine. 2020;12(1):68.

31. Lau SY, Wang P, Mok BW, et al. Attenuated SARS-CoV-2 variants with deletions at the S1/S2 junction. Emerg Microbes Infect. 2020;9(1):837–842.

32. Liu Z, Zheng H, Lin H, et al. Identification of Common Deletions in the Spike Protein of Severe Acute Respiratory Syndrome Coronavirus 2. J Virol. 2020;94(17).

33. Ogando NS, Dalebout TJ, Zevenhoven-Dobbe JC, et al. SARS-coronavirus-2 replication in Vero E6 cells: replication kinetics, rapid adaptation and cytopathology. J Gen Virol. 2020;101(9):925–940.

34. Tyrrell DA, Bynoe ML. Cultivation of a novel type of common-cold virus in organ cultures. Br Med J. 1965;1(5448):1467–1470.

35. Holwerda M, Kelly J, Laloli L, et al. Determining the Replication Kinetics and Cellular Tropism of Influenza D Virus on Primary Well-Differentiated Human Airway Epithelial Cells. Viruses. 2019;11(4):377.

36. Kendall EJ, Bynoe ML, Tyrrell DA. Virus isolations from common colds occurring in a residential school. Br Med J. 1962;2(5297):82–86.

37. Laporte M, Stevaert A, Raeymaekers V, et al. The SARS-CoV-2 and other human coronavirus spike proteins are fine-tuned towards temperature and proteases of the human airways. bioRxiv. 2020:2020.2011.2009.374603.

38. Velthuis AJ. Common and unique features of viral RNA-dependent polymerases. Cell Mol Life Sci. 2014;71(22):4403–4420.

39. Mackman RL, Hui HC, Perron M, et al. Prodrugs of a 1′-CN-4-Aza-7,9-dideazaadenosine C-Nucleoside Leading to the Discovery of Remdesivir (GS-5734) as a Potent Inhibitor of Respiratory Syncytial Virus with Efficacy in the African Green Monkey Model of RSV. Journal of Medicinal Chemistry. 2021;64(8):5001–5017.

40. Vanderheiden A, Ralfs P, Chirkova T, et al. Type I and Type III Interferons Restrict SARS-CoV-2 Infection of Human Airway Epithelial Cultures. J Virol. 2020;94(19).

